# Rapid-kinetics degron benchmarking reveals off-target activities and mixed agonism-antagonism of MYB inhibitors

**DOI:** 10.1101/2023.04.07.536032

**Authors:** Taku Harada, Monika W. Perez, Jérémie Kalfon, Flora Dievenich Braes, Rashad Batley, Kenneth Eagle, Behnam Nabet, Becky Leifer, Jasmin Kruell, Vikram R. Paralkar, Kimberly Stegmaier, Angela N. Koehler, Stuart H. Orkin, Maxim Pimkin

## Abstract

Attenuating aberrant transcriptional circuits holds great promise for the treatment of numerous diseases, including cancer. However, development of transcriptional inhibitors is hampered by the lack of a generally accepted functional cellular readout to characterize their target specificity and on-target activity. We benchmarked the direct gene-regulatory signatures of six agents reported as inhibitors of the oncogenic transcription factor MYB against targeted MYB degradation in a nascent transcriptomics assay. The inhibitors demonstrated partial specificity for MYB target genes but displayed significant off-target activity. Unexpectedly, the inhibitors displayed bimodal on-target effects, acting as mixed agonists-antagonists. Our data uncover unforeseen agonist effects of small molecules originally developed as TF inhibitors and argue that rapid-kinetics benchmarking against degron models should be used for functional characterization of transcriptional modulators.

## Introduction

Transcription factors (TFs) establish cell states by directly interpreting the cis-regulatory code of the genome.^1,2^ TF dysregulation by mutation or aberrant expression underlies numerous diseases, and is a hallmark of cancer.^3,4^ In principle, the highly specific roles of TFs in enforcing developmental and disease phenotypes make them ideal targets for drug development.^5–7^ However, direct drugging of TFs remains challenging despite extensive efforts.^5,6^ A central problem in these efforts is a lack of detailed mechanistic understanding of TF function and, as a result, no clear consensus on how the functional output of a TF should be measured for drug characterization purposes.^5^ Recently, direct gene-regulatory functions of TFs have been established in pre-steady state assays where rapid TF degradation is coupled with measurements of genome-wide transcription rates.^8–13^ These studies have demonstrated that TFs have narrow direct transcriptional programs and that long-term TF deprivation (e.g. after a CRISPR/Cas9-mediated knockout) leads to significant secondary effects obscuring a direct functional readout.^8,11^

We reasoned that the specificity and on-target activity of TF inhibitors would be best evaluated in a rapid kinetics system where their immediate transcriptional effects are benchmarked against targeted TF degradation (Figure 1A).

**Figure 1.**
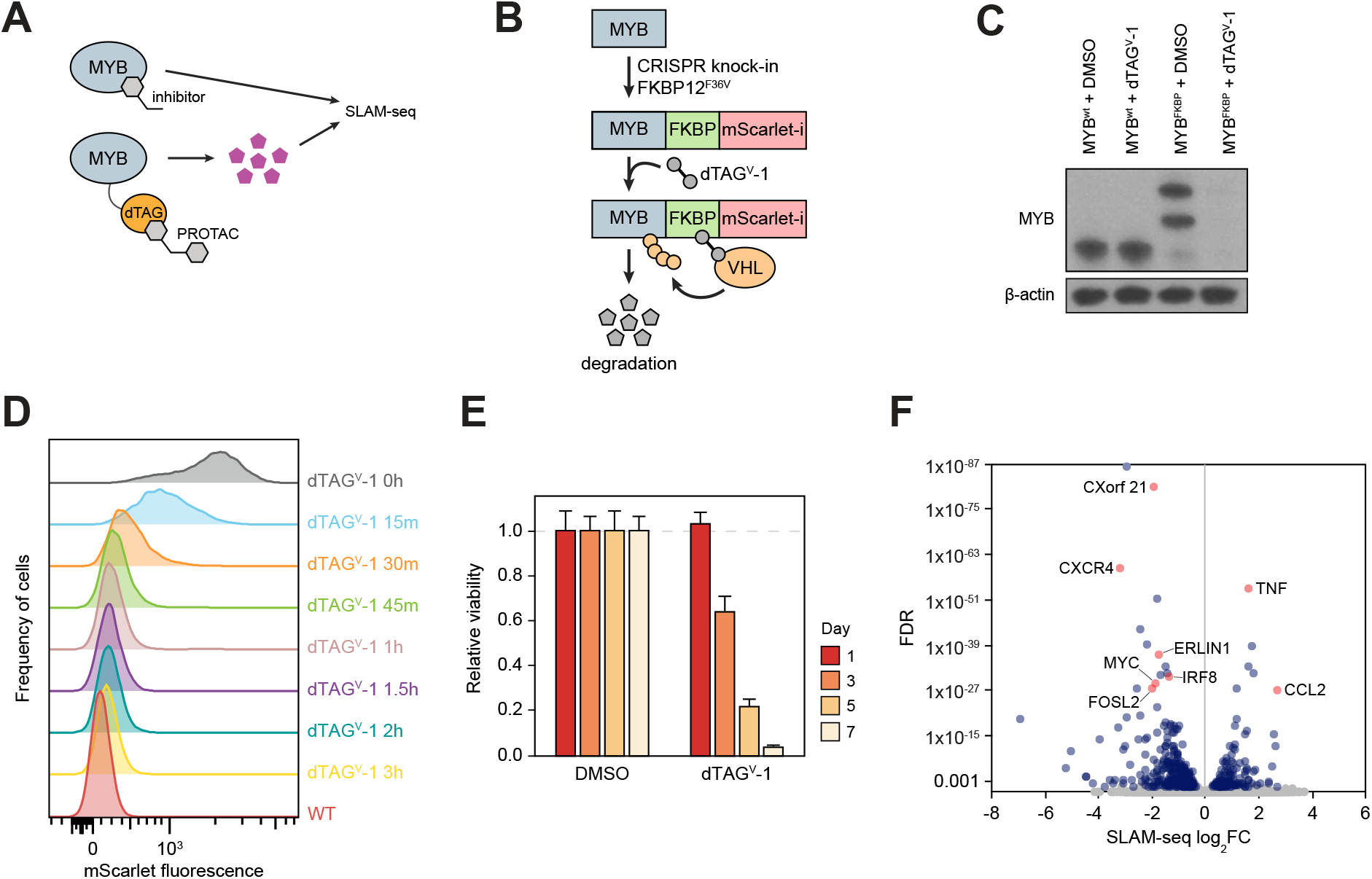
A chemical degron model of MYB. (A) Study overview. (B and C) Schematic (B) and Western blot (C) demonstrating endogenous MYB degron tagging by CRISPR-HDR and subsequent targeted degradation of the fusion protein. (D) A time course experiment where MYB degradation is followed by FACS measurement of the fusion protein fluorescence. (E) Relative cell viability measured at various points after MYB degradation. (F) Volcano plot of genome-wide changes in transcription rates measured by SLAM-seq 1 hour after MYB degradation.

In particular, since expression of many TFs rapidly changes in response to various external stimuli^3,5,14–16^, a rapid-kinetics assay is necessary to distinguish the direct effects of an inhibitor on its target TF from the secondary effects resulting from a potential change in expression of the target TF. Focusing on MYB, an oncogenic TF driver of multiple cancers^17–24^ and an emerging therapeutic target^5,25–27^, we engineered a chemical degron model and established the direct gene-regulatory functions of MYB in a nascent transcriptomics assay. By benchmarking the nascent transcriptomics signatures of six MYB inhibitors against the degron we uncovered their off-target effects and unexpected mixed agonism-antagonism of their on-target activities.

## Results

### A degron model reveals direct gene-regulatory functions of MYB

MYB is a critical transcriptional dependency of acute myeloid leukemia (AML)^11,18,28,29^, where it has been a long-term focus of therapeutic efforts^30–37^, making AML a relevant context for functional characterization of MYB inhibitors. We therefore began by engineering a chemical degron model of MYB in AML cells. We fused the C-terminus of MYB with an FKBP12^F36V^ (dTAG) domain and a fluorescent tag by a homozygous knock-in of the FKBP12^F36V^-mScarlet-coding DNA sequence into the endogenous MYB locus in MV411 cells (Figure 1B,C). The resulting fusion protein was nearly completely degraded after a 1-hour treatment with dTAG^V^-1, a highly specific VHL-engaging PROTAC^38^ (Figure 1D). As expected, degradation of MYB resulted in a profound loss of cell viability, consistent with the effects of a genetic MYB knockout in AML cells^11^ (Figure 1E). To establish MYB’s direct gene-regulatory functions, we measured genome-wide rates of nascent mRNA synthesis by thiol (SH)-linked alkylation metabolic sequencing of RNA (SLAM-seq)^39^ after a 1-hour MYB degradation. Defining direct targets as those genes which displayed significant changes in transcription rates (FDR<0.05)^8^, we detected 450 genes directly regulated by MYB (Figure 1F). Of these, 319 genes were downregulated and 131 genes were upregulated, indicating that MYB acts as a transcriptional activator and repressor of these genes, respectively.

### Degron benchmarking establishes target specificity of MYB inhibitors

In parallel, we performed SLAM-seq in AML cells treated with six agents reported as MYB inhibitors: MYBMIM^31^, celastrol^34^, naphthol AS-E phosphate^40^, mebendazole^32,41,42^, plumbagin^33^ and all-trans retinoic acid (ATRA)^43^ (Figure 1A, Table 1). Although each is described as an inhibitor of MYB, the agents act through different mechanisms.

**Table 1.** Inhibitors used in the study.

| Inhibitor | Mechanism | IC50 in MV411, μM | Concentration used, μM | References |
| --- | --- | --- | --- | --- |
| ATRA | ↓MYB expression | 0.27 | 1 | 43 |
| Celastrol | MYB::p300 (KIX domain) | 0.12 | 0.1 | 34 |
| Mebendazole | MYB degradation | 0.71 | 10 | 41,42,62 |
| MYBMIM | MYB::CBP/p300 (KIX domain) | 1.5 | 20 | 31 |
| Naphthol AS-E phosphate | MYB::p300 (KIX domain) | 13.2 | 10 | 40 |
| Plumbagin | MYB::p300 (MYB TAD) | 0.58 | 0.5 | 33 |
| JQ1 | BET bromodomain inhibition → ↓MYB functional output | 0.41 | 1 | 44,45 |
| KI-MS2-008 | Stabilization of MAX homodimer → MYC inhibition | 22.4 | 50 | 46 |
| MYCi361 | Dissociation of MYC/MAX heterodimer → MYC inhibition | 2.22 | 10 | 47 |

MYBMIM, celastrol, plumbagin and naphthol AS-E phosphate inhibit MYB function by disrupting its interaction with the critical co-activator p300 and are therefore expected to cause an immediate dysregulation of the MYB target genes. In contrast, ATRA acts indirectly by decreasing MYB expression and thus its immediate transcriptional effects are expected to be distinct from the effects of acute MYB loss. Although the direct target of mebendazole has not been identified, it appears to target MYB for proteolytic degradation with prolonged exposure and may act through additional mechanisms.^42^ For comparison, we treated MV411 cells with the BET bromodomain inhibitor JQ1^44^, which has been reported to indirectly inhibit MYB by interfering with its expression and function^45^. In addition, given that MYB appears to directly activate the expression of MYC^28^ (Figure 1F), we included in the comparison two MYC inhibitors, KI-MS2-008^46^ and MYCi361^47^. MYBMIM, a cell-permeable peptidomimetic^31^, was applied for 30 minutes while all other inhibitors were applied for 1 hour prior to SLAM-seq.

The MYB inhibitors displayed a dramatic variability in the number of dysregulated genes, varying from 19 (naphthol) to 1123 (MYBMIM; Figure 2A). Consistent with a prior report^8^, JQ1 caused widespread and bimodal effects, altering the transcription rates of >2000 genes in both directions. On pairwise overlap, the MYB inhibitors captured a relatively minor portion of the direct MYB program (between 5-155, or 1-34%, of the 450 direct MYB targets), compared with 43% of the MYB program captured by JQ1 (Figure 2B). Nonetheless, the MYB inhibitors displayed a stronger specificity for MYB target genes compared to JQ1, because they elicited much narrower responses (Figure 2C). Surprisingly, ATRA, which, as expected^43^, directly inhibited MYB transcription, also demonstrated a strong enrichment for primary MYB targets (Figure 2C), perhaps due to cooperation between MYB and the retinoid receptors at the target gene level^48^. The MYC inhibitors were also enriched for the MYB targets but displayed extremely narrow programs (Figure 2A,C). Overall, approximately half of the direct MYB program (249 of 450, or 55% target genes) was captured by any combination of MYB inhibitors, while 86 targets were affected by 2 or more MYB inhibitors (Figure 2D,E). We considered the possibility that, because the definition of a primary MYB target depends on the significance cutoff, some genomic effects of the MYB inhibitors may be falsely classified as off-target if the same genes fell just below the significance level in the degron SLAM-seq. We found 1610 genes whose transcription rates were affected by at least one MYB inhibitor but were unchanged after MYB degradation (Figure 2D). A significant majority of these genes (1171, 72%) displayed less than a 25% net change in the transcription rate after a near-complete MYB degradation (Figure 2F), thus likely representing bona-fide off-target effects of the MYB inhibitors. We conclude that, while MYB inhibitors display strong enrichments for primary MYB targets, they do not attenuate the entire MYB transcriptional program, and significant portions of their activities appear to be off-target.

**Figure 2.**
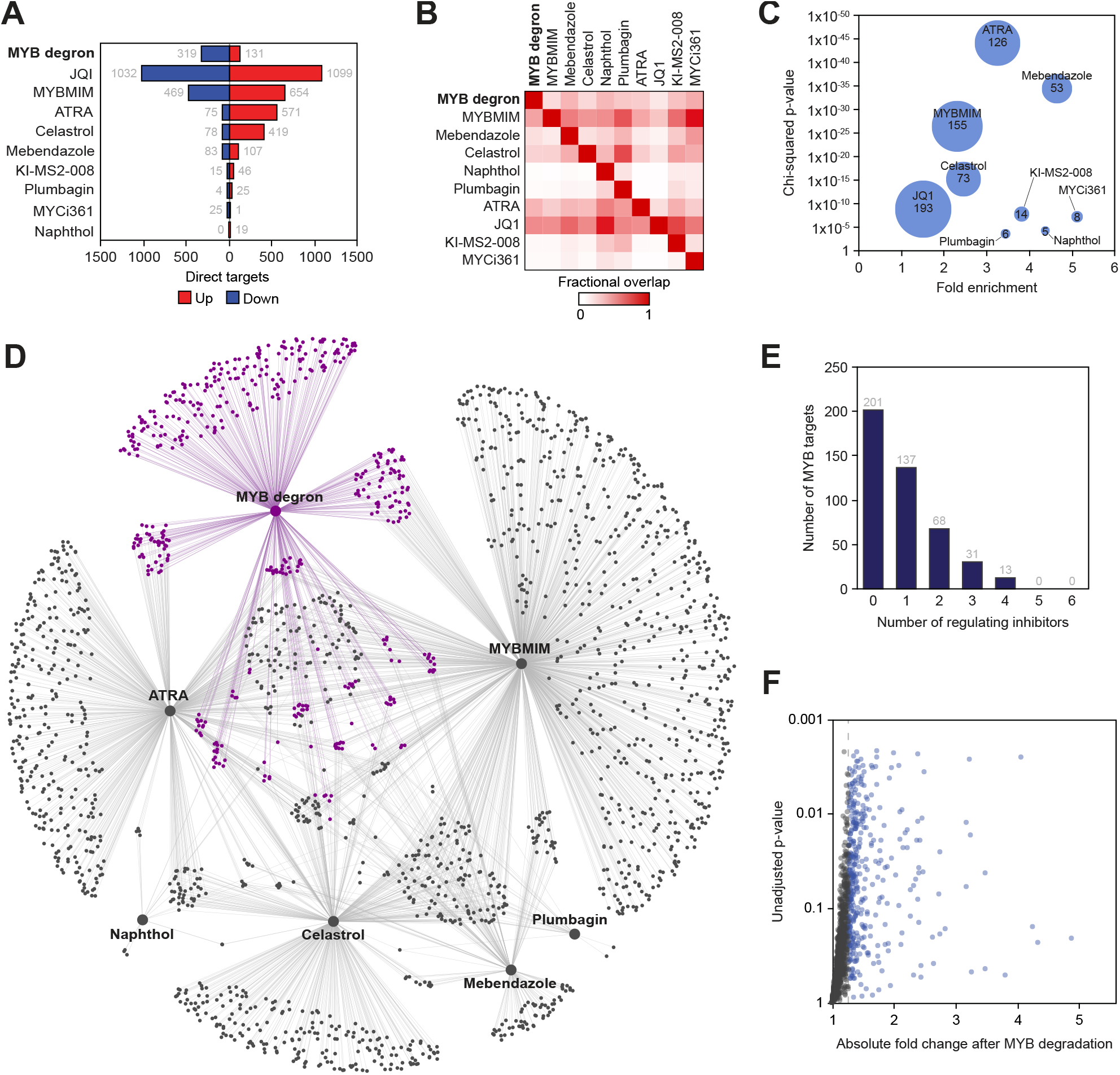
Direct gene-regulatory programs of MYB and its inhibitors. (A) Numbers of direct targets of MYB and transcriptional inhibitors visualized according to the direction of response to MYB degradation or inhibitor treatment. (B) Fractional pairwise overlap between the direct transcriptional targets of MYB and inhibitors. Data points reflect fraction of columns overlapping with rows. (C) Enrichment of MYB targets among the direct targets of transcriptional inhibitors. Bubble size represents the number of overlapping genes. (D) Visualization of the reconstructed regulatory network, where large nodes represent MYB degron and inhibitors, small nodes represent target genes and edges represent direct regulatory connections. MYB targets and edges are highlighted in purple. (E) Overlap of the MYB inhibitors among the direct MYB targets, visualized as number of direct MYB targets directly regulated by *n* inhibitors. (F) Genes whose transcription rates were affected (SLAM-seq FDR <0.05) by at least one MYB inhibitor but unaffected by targeted MYB degradation (n = 1610) are depicted according to the absolute fold change and unadjusted p-value in the SLAM-seq assay performed after MYB degradation.

### MYB inhibitors act as mixed agonists-antagonists

Having established target specificities of the MYB inhibitors, we sought to characterize their on-target functional outputs. Contrary to the effects of MYB degradation, the MYB inhibitors generally activated more genes than they repressed (Figure 2A). In addition, we observed generally weak correlations between the MYB degron and inhibitor responses, both transcriptome-wide and in the space of the confirmed direct MYB targets (Figure 3A,B). Plotting the transcriptional responses to MYB inhibitors against the degron-induced changes at overlapping target genes revealed that the inhibitors displayed bimodal effects on the transcription of MYB-regulated genes, further activating subsets of genes that were repressed by MYB degradation, and vice versa (Figure 3C). We further reasoned that the bimodal activities of the inhibitors may be distributed unevenly across ontologically defined groups of inhibitor-responsive genes. Indeed, utilizing comparative pathway enrichment analysis we detected distinct MYB-regulated pathways where the action of MYB was either agonized or antagonized by the inhibitors (Figure 3D,E). We conclude that MYB inhibitors augment, rather than reverse, the gene-regulatory functions of MYB at a subset of MYB-regulated genes and pathways, thus acting as mixed agonists-antagonists.

**Figure 3.**
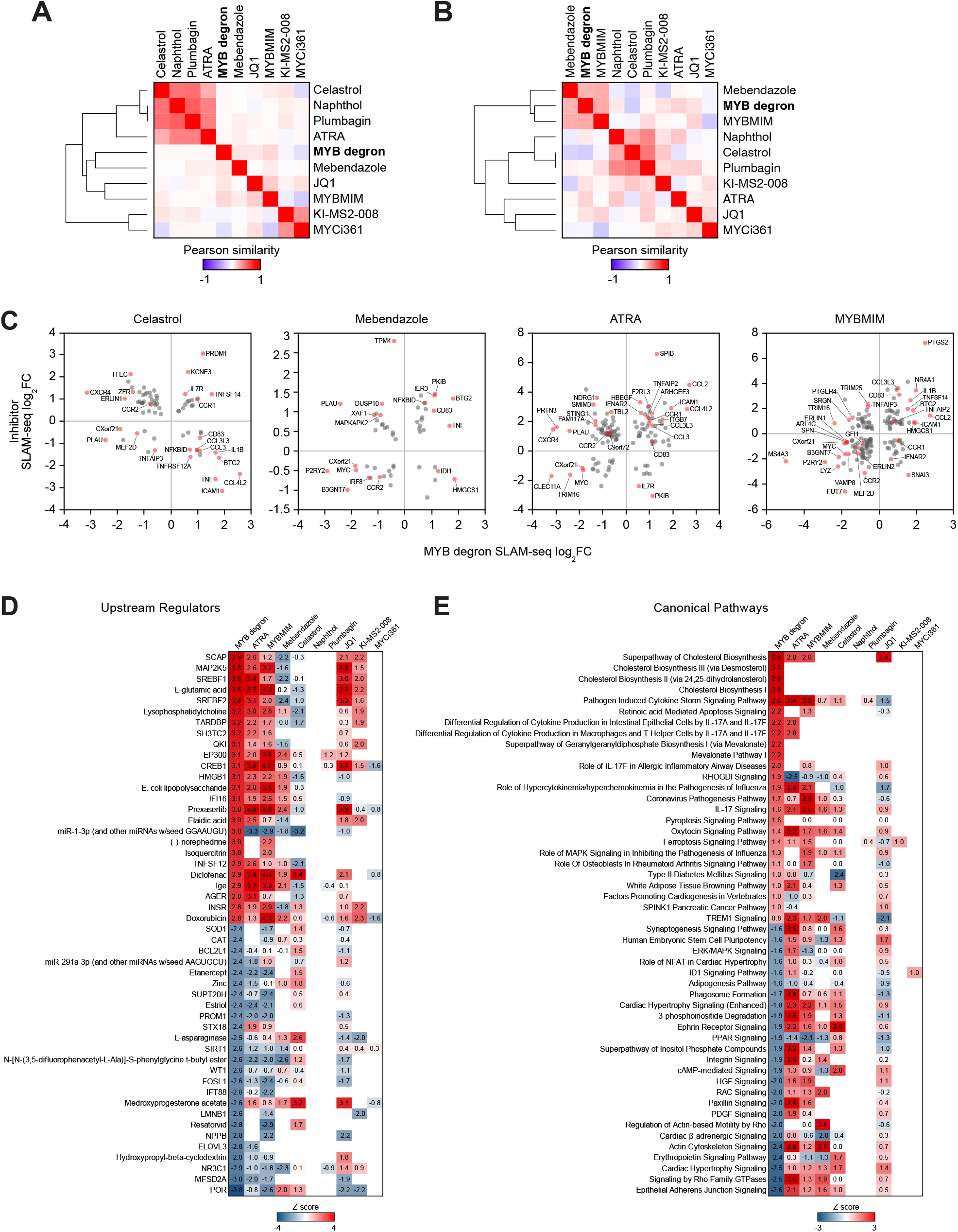
On-target activities of MYB inhibitors. (A and B) Pairwise Pearson similarity of nascent transcriptional responses to MYB degradation and inhibitor treatment in the space of all expressed genes (A; n=7485) and direct MYB targets (B; n=450), organized by unsupervised hierarchical clustering with complete linkage. (C) SLAM-seq responses of MYB target genes to MYB degradation vs. inhibitor treatment, demonstrating mixed agonist-antagonist effects of MYB inhibitors on the transcription of MYB target genes. (D and E) Heatmaps of predicted activation scores (z-scores) of top enriched upstream regulator (D) and canonical (E) pathways among the genes affected by MYB degradation and inhibitor treatment. Fifty pathways with top z-scores in the MYB degron dataset are visualized. Data points reflect pathways with significant enrichments (BH-adjusted *p*-value <0.05, calculated by internal IPA function) and are colored according to the activity z-scores, predicting pathway inhibition (z-score <1) or activation (z-score >1).

## Discussion

TFs control gene expression through complex interactions with chromatin, which, in addition to DNA, includes numerous cooperating TFs, cofactors and structural proteins^1,2^. The molecular mechanisms linking TF binding to DNA to a change in gene expression remain poorly understood, but it is clear that a TF’s functional output depends exquisitely on the local chromatin context^2,8,9,11^. Although TFs have been historically characterized as activators or repressors, many function as both, depending on the local presence of cooperating factors^46,49,50^. The combinatorial nature of these interactions and absence of simple rules governing TF output significantly complicates evaluation of small-molecule inhibitors and makes elusive such fundamental pharmacodynamic parameters as specificity and efficacy. Indeed, the functional effect of a hypothetical small-molecule TF modulator may vary dramatically across the expressed genome even if it does not engage in any off-target interactions. In addition, chemical TF inhibition may not be functionally equivalent to TF loss, depending on the structural role a TF may play in shaping the local chromatin environment (for example, competing with other TFs for DNA binding)^51^. We illustrate these concepts by a comparative analysis of six MYB inhibitors versus chemical MYB degradation, where we use the degron as a proxy model of MYB inhibition with near-absolute specificity and efficacy.

Although all of the evaluated MYB modulators demonstrated some effects on primary MYB targets, acting with more specificity than BET bromodomain inhibition, they were unable to capture the entire MYB program and displayed significant off-target activities. We speculate that this apparent lack of specificity may be partly driven by their indirect actions on MYB and the complexity of the MYB and p300/CBP interactomes, rather than simply by engagement of unintended targets or poor on-target efficacy. Indeed, four MYB inhibitors disrupt the interaction between MYB and the co-activators p300/CBP (celastrol, plumbagin, MYBMIM and naphthol AS-E phosphate; Table 1). Of these, plumbagin appears to bind the trans-activator domain of MYB^33^, while the other three inhibitors bind to the KIX domain on the surface of p300/CBP and thus modulate MYB activity indirectly^31,34,40^. The KIX domain of p300/CBP interacts with a number of other TFs^52^, and therefore the “off-target” effects of the KIX-binding inhibitors may in fact represent on-target modulation of p300/CBP in a MYB-independent manner. Conversely, while the interaction with p300/CBP is essential for the oncogenic properties of MYB^53,54^, it does not represent the full repertoire of MYB interactions^22,30,55,56^, and any p300/CBP-independent activities of MYB will be preserved after p300/CBP inhibition.

Functional agonism, antagonism and mixed agonism-antagonism are basic parameters initially developed in receptor pharmacology where they are typically established by comparing drugs with natural receptor ligands.^57^ While TF inhibitors are typically developed as antagonists, in principle they can augment the local output of a TF, whether it be to activate or repress the transcription of a target gene. For example, some inhibitors of MYC act by stabilizing MAX homodimers on DNA and can be thought of as MAX agonists^46^. Our data uncover unexpected activities of MYB inhibitors as context-dependent agonists. Although p300 and CBP are typically classified as coactivators, they may also function as corepressors^58,59^. We speculate that p300/CBP restrain MYB function at a subset of target genes and inhibiting their interaction with MYB may result in increased MYB activity. Surprisingly, ATRA, which regulates gene expression through binding to the nuclear retinoid receptor^60,61^, demonstrated a strong enrichment for primary MYB targets and similar functional ambivalence, suggesting a potential interaction between MYB and the retinoid receptor at the enhancer/promoter level. Importantly, the mixed agonism-antagonism of MYB modulators differentially affects distinct MYB-regulated pathways, indicating that any therapeutic use of these molecules would need to be tailored to the specific parts of the MYB program that drive the disease phenotype.

In conclusion, we demonstrate that the rapid kinetic resolution of TF degron models allows for a more precise characterization of the target specificity and efficacy of TF-directed inhibitors. Our observations indicate that benchmarking of TF modulators against degron models in nascent transcriptomics assays should be considered as an important criterion in their functional characterization.

## Methods

### Cell lines

MV411 cell lines were cultured in the RPMI-1640 media containing 10% fetal bovine serum and regularly tested to be free of *Mycoplasma spp*.

### Western blotting

Whole-cell lysates were prepared in RIPA buffer (Boston Bio-Products BP-115-500) with protease inhibitor cocktail (ThermoFisher 23225). Lysates were boiled in Laemmli buffer (BioRad 1610737), separated by SDS-PAGE, and transferred and blocked using standard methodology. HRP-conjugated anti-mouse and anti-rabbit IgG secondary antibodies were used for imaging (BioRad 1706515 and 1706515) with an enhanced chemiluminescence substrate (PerkinElmer NEL104001EA) according to manufacturers’ instructions.

### Targeted TF degradation

MV411 cells were modified by CRISPR-HDR to express a C-terminal FKBP12^F36V^ (dTAG) fusion of MYB. A donor DNA construct, including the knock-in cassette and ca. 400 homology arms, was commercially synthesized (Genewiz, Burlington, MA) and cloned into the pAAV-MCS2 plasmid vector obtained from Addgene (Watertown, MA). rAAV packaging was performed at the Boston Children’s Hospital Viral Core. MV411 cells were electroporated with Cas9/sgRNA complexes targeting the HDR insertion using a Lonza SF Cell Line 4D Nucleofector (Lonza V4XC-2032). RNP complexes were formed by mixing 8.5 μg of TrueCut Cas9 Protein v2 (Invitrogen A36499) and 120 pmol sgRNA. 0.3×10^6^ cells were washed with PBS and resuspended in 20 μL of SF Cell Line solution (Lonza). Ten μL of crude rAAV lysate was added to the cells immediately after electroporation^63^. After a 5-7 day incubation period the cells were sorted for mScarlet fluorescence. Single clones were then obtained by single-cell dilution microwell plating and screened for bi-allelic donor insertion by PCR. Clones were validated by Western blotting and Sanger sequencing. TF degradation was induced by adding 500 nM of dTAG^v^-1 as previously described^38^ and followed by FACS measurement of mScarlet fluorescence and Western blotting.

### SLAM-seq

For thiol (SH)-linked alkylation metabolic sequencing of RNA (SLAM-seq)^39^ 2.5×10^6^ MV411 cells per replicate were incubated with 500 nM dTAG^V^-1, or DMSO, for 1 hour. For inhibitor experiments, MV411 cells were treated with indicated concentrations of inhibitors (Table 1) for 30 min (MYBMIM) or 1 hour (all other inhibitors). All experiments were done in at least 4 replicates. Metabolic labeling was performed by adding S^4^U to a final concentration of 100 μM for an additional hour. Cells were flash-frozen and total RNA was extracted using Quick-RNA MiniPrep (Zymo Research) according to the manufacturer’s instructions except including 0.1 mM DTT to all buffers. Thiol modification was performed by 10 mM iodoacetamide treatment followed by quenching with 20 mM DTT. RNA was purified by ethanol precipitation and mRNA-seq was performed as described above. A modified version of the slamdunk pipeline was used for SLAM-seq processing (available at https://github.com/jkobject/slamdunk).

## Acknowledgements

This work was supported by research grants to MP, SHO and ANK from the Adenoid Cystic Carcinoma Research Foundation. KS was supported by NIH 5R35 CA210030 and NIH P50 CA206963. BN was supported by NCI K22 CA258805. We thank Dr. A. Kentsis for providing MYBMIM.

## Author Contributions

Conceptualization: SHO, MP, ANK, KS. Data analysis: MWP, JK, MP, KE, BL, VRP, KS, ANK. Experiments: TH, MP, BN, FDB, RB. Visualization: FDB, MWP, MP. Funding acquisition: SHO, MP. Supervision: SHO, MP. Writing of the manuscript: MP and SHO with input from all authors.

